# Evolving Threats: Leveraging *C. elegans* to Decode the Virulence Profiles of Highly Related Environmental *Salmonella* Newport Isolates

**DOI:** 10.1101/2025.09.29.679144

**Authors:** Christina M. Ferreira, Mirae Choe, Bella Wayhs, Julie Haendiges, Robert Literman, Jianghong Meng, Arjuman Ghazi, Rebecca L. Bell

## Abstract

*Salmonella enterica* subspecies *enterica*, particularly serovar Newport, remains a leading cause of foodborne illnesses in the United States, implicated in numerous outbreaks associated with a diverse array of food products. This study rigorously investigates the virulence of five distinct *S*. Newport isolates, characterized by varying patterns of pulse-field gel electrophoresis (PFGE) molecular-diagnostic subtyping, using the nematode *Caenorhabditis elegans* as a host model organism. We conducted viability assays on *C. elegans* to evaluate how these isolates affect nematode survival. The selected bacterial strains, rooted in historical foodborne outbreak significance but environmentally isolated, were previously sequenced to provide a comprehensive genomic framework. A notable focus of our research was on the nearly genetically identical PFGE types Newport-61 and the Newport-1015 isolates, which differ by a ∼1.7 Mb genomic inversion. *C. elegans* survival assays in response to pathogenic-strain infections revealed that one Newport-1015 and the Newport-61 were particularly more virulent compared to other strains tested. These findings enhance our understanding of the pathogenic potential of environmental *S*. Newport and highlight the need to understand the regulatory mechanisms that contribute to virulence capacity.

## Introduction

*Salmonella enterica* subspecies *enterica* is the leading cause of microbial foodborne illnesses in the United States, resulting in over 1 million infections and more than 400 deaths each year (1, 2). Among its serovars, *S. enterica* serovar Newport (Newport) ranks as the second most common culture-confirmed serovar associated with foodborne outbreaks in the country. Newport has a broad host range, implicated in illnesses linked to a variety of foods, including tomatoes, cucumbers, papaya, onions, and ground beef (3-8). From 2000 to 2020, there were 227 foodborne salmonellosis outbreaks attributed to Newport, with six of those being recurrent outbreaks occurring over a 12-year period connected to tomatoes from the Virginia Eastern Shore (VES) (3, 9, 10). These tomato outbreaks were linked to a clonal strain of Newport, identified through PFGE as “Pattern 61” (XbaI JJPX01.0061, Newport-61). This clone is now categorized by the CDC under REPJJP03, which also includes Newport Pattern 1015 (XbaI JJPX01.1015, Newport-1015).

The nematode *Caenorhabditis elegans* has been established as a valuable model organism for studying the virulence mechanisms of human pathogens. Previously, worms exposed to *S. enterica* serovar Typhimurium (Typhimurium) strains were shown to have a significantly shortened lifespan with visible signs of infection in the intestinal tract. Notably, *Salmonella* strains that exhibit reduced virulence in mammals were shown to have similarly attenuated impact on the lifespan of *C. elegans* (11, 12). This indicates that bacterial genes important for pathogenic potency in vertebrate hosts are likely similarly required during *C. elegans* infection, highlighting its value to assess pathogenic virulence and host-pathogen interactions.

Newport-61 and Newport-1015 have been isolated from the surface water and sediment of the VES, with a seasonal recurrence of clonal isolates (13, 14). However, there are few documented cases, prior to the retirement of CDC PFGE PulseNet, of Newport-1015 causing human infection, suggesting overall less virulence capacity. In this study, we used *C. elegans* to evaluate the virulence of five *S*. Newport environmental isolates, each representing different PFGE patterns of historical significance related to foods. These isolates are closely related, with Newport-1015 and Newport-61 being particularly noteworthy, as they are nearly genetically identical except for a ∼1.7 Mb inversion (15).

## Materials and Methods

### Bacterial Isolates

Strains used in the *C. elegans* experiment included *E. coli* OP50 (negative control) and six *Salmonella enterica* strains found in Table 1. The Typhimurium isolate (CFSAN000741) was used as the positive control for the *C. elegans* assays, and the Newport isolates (CFSAN000859, CFSAN001461, CFSAN001891, CFSAN003353, and CFSAN001469) were previously reported (15). All strains were maintained as frozen stocks in Brain Heart Infusion Broth (BHIB) with 50% glycerol (v/v), and experiments were conducted with freshly grown cultures, streaked from frozen stock cultures onto Luria Bertani agar and incubated at 35°C ± 2°C for 22 ± 2 hours (16). Single colonies from these plates were inoculated into 3mL of Luria Bertani broth (LB) and incubated statically at 35°C ± 2°C for 22 ± 2 hours.

**Table 1:** *Salmonella* strains used for bioinformatic analysis and/or *C. elegans* assays.

| Sample ID | Serovar | Source | Sequence Type | PFGE Pattern | Genome Accession Number |
| --- | --- | --- | --- | --- | --- |
| CFSAN001891 | Newport | Goose feces | 118 | JJPX01.1015 | CP140739 <sup>(15)</sup> |
| CFSAN001461 | Newport | Sediment | 118 | JJPX01.1015 | CP140738 <sup>(15)</sup> |
| CFSAN003353 | Newport | Sediment | 118 | JJPX01.0061 | CP140737 <sup>(15)</sup> |
| CFSAN001469 | Newport | Sediment | 350 | JJPX01.0044 | CP140735 <sup>(15)</sup> |
| CFSAN000859 | Newport | Stool | 5 | JJPX01.0011 | CP140750 <sup>(15)</sup> |
| CFSAN000741 | Typhimurium | Tissue | -- | -- | JBREBA000000000 |

### WGS Comparative Analysis

Strains used in this study were previously sequenced to provide complete, reference quality genomes using the PacBio (17). The assemblies were uploaded to NCBI and annotated using the NCBI Prokaryotic Genome Annotation Pipeline (PGAP) (18). Roary was used to generate a pangenome alignment and to identify genes that were unique to the different PFGE patterns (19). Visualization of the output from Roary were created using Phandango (20). AMRFinderPlus, Phastest, and SPIFinder were used to identify AMR genes, phages, and pathogenicity islands (21-23). BLAST was used to identify the percent identity of the strains (24). Geneious Prime (v 2025.1.2) was used for *in silico* restriction digestion of the strains with the enzyme XbaI to identify the difference in PFGE pattern comparison of JJPX01.0061 and JJPX01.1015.

### *Caenorhabditis elegans* Survival Assay

*C. elegans* survival experiments were conducted with the wild-type strain, N2. Worms were maintained using standard techniques on nematode growth medium (25) plates seeded with normal laboratory diet of *Escherichia coli* strain OP50 (OP50) at 20°C. Fresh NGM plates were seeded with 70μL of OP50 or one of the six *Salmonella* pathogenic strains. OP50 plates were allowed to dry for 24 hours before adding worms. *Salmonella*-seeded plates were used after 5 hours of drying; hence fresh *Salmonella* plates were prepared every 24 hours.

For the survival assays, healthy, gravid young adult worms were transferred to fresh OP50-seeded NGM plates, allowed to lay eggs for 1–2 days and removed. The eggs were reared at 20°C and age-matched L4-stage, pre-adult larvae were selected for the experiment and transferred to NGM plates seeded with either the OP50 control bacteria or various *Salmonella* strains. Thirty L4-stage worms were transferred to each plate, with a total of 150 worms tested per bacterial strain. Worm survival was scored twice daily by gently touching the head, tail, or midsection with a platinum wire pick; worms were considered dead if they failed to respond.

Worms were transferred to fresh, corresponding plates every 24 hours. Survival monitoring continued until all original worms died, using 12-hour/12-hour or 8-hour/16-hour scoring intervals. Worms that exploded, bagged, crawled off the plate, or were otherwise unaccounted for were censored from analysis. All survival assays were conducted twice in two biological replicates. Survival statistics were plotted using the Kaplan–Meier method. Statistics were calculated using the nonparametric log-rank Mantel−Cox method on the OASIS2 platform and subjected to multiplicity Bonferroni correction (26).

## Results and Discussion

### Novel inversion in highly related S. Newport isolates suggests host specific adaptation

The strains presented in this study are all known to cause human infections to varying degrees and have been identified in across the United States. Gene content analysis shows that these isolates demonstrate over 98% genetic identity to one another, with Newport-61 being more than 99% identical to Newport-1015 (Figure 1). It was previously reported that two strains, CFSAN001891 and CFSAN001461 (both JJPX01.1015), have an approximately 1.7 Mbp inversion relative to the chromosome of CFSAN003353 (JJPX01.0061) (15). As can be seen in Figure 1, the 1.7 Mbp inversion account for the difference in PFGE patterns between JJPX01.0061 and JJPX01.1015 as the location of the XbaI restriction enzyme site is altered (Figure 2, indicated by “X”).

**Fig 1.**
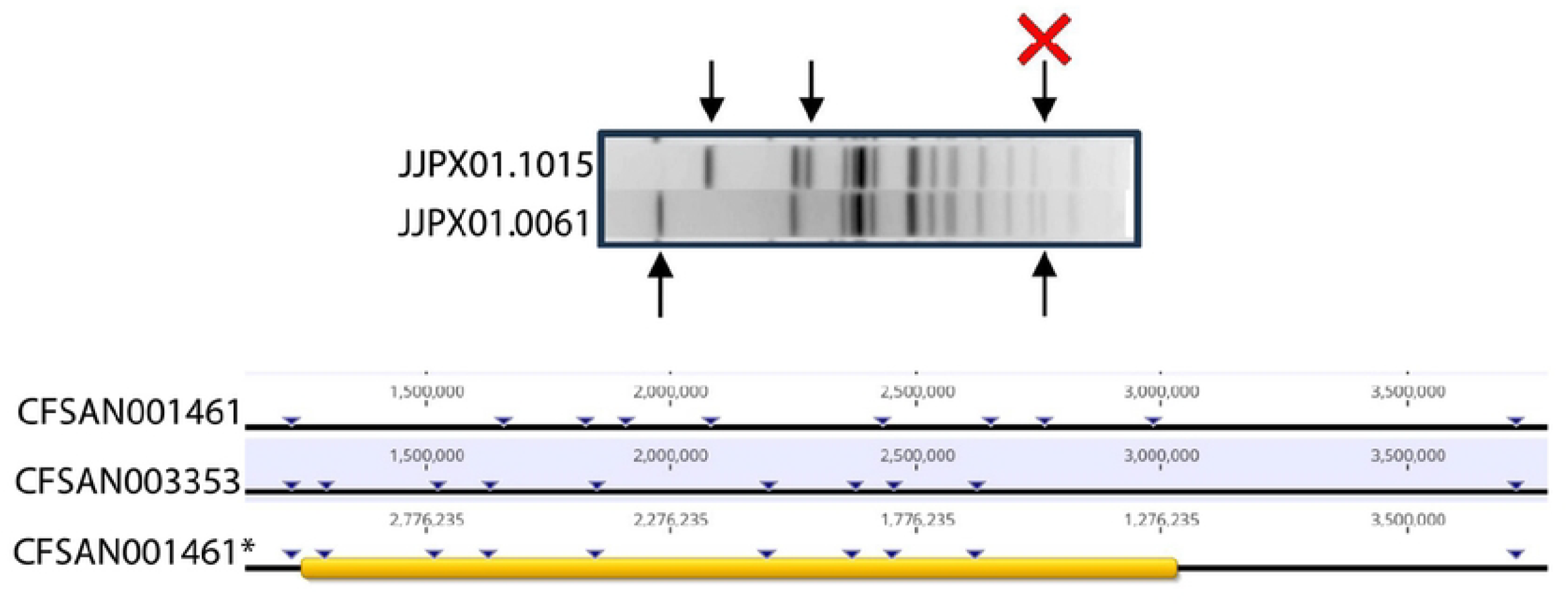
Phylogeny showing gene presence/absence of key *Salmonella* Newport isolates. The phylogenetic tree shows the relatedness of the *S*. Newport isolates to one another. The blue colored area represents the presence/absence of genes across the entirety of the genomes. The sequence types represented include 118 (magenta), 350 (yellow) and 5 (dark blue). Analysis of genetic content related to virulence, AMR or prophages did not show any differences among the isolates in this study (data not shown).

**Fig 2.**
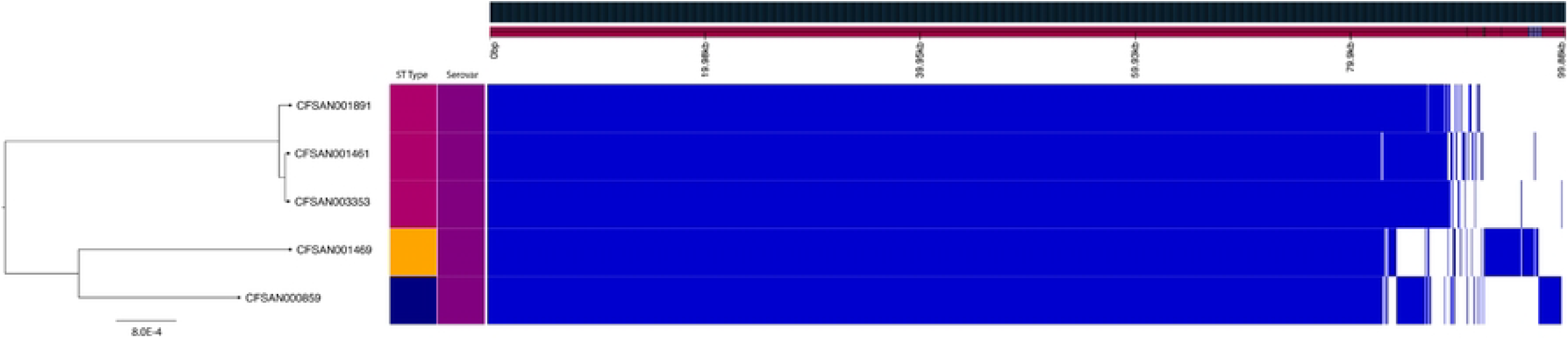
Inversion confirmation of CFSAN001461 and CFSAN1891 using *in silico* PFGE analysis. PFGE XbaI pattern of JJPX01.1015 compared to JJPX01.0061 (top), where the arrows highlight the differing bands in the PFGE patterns and the red “X” denotes the band that is not present in the 1015. The inversion identified in JJPX01.1015 accounts for the differences in the PFGE patterns due to changes in cut site locations within the genome. The locations of these cut site changes (bottom) due to the directionality of the inversion - by manually rotating the 1.7Mbp JJPX01.1015 inversion at the Gifsy-1 sites (denoted by CFSAN001461*) and subsequently performing *in silico* digestion with XbaI, the cut sites revert to the same fragment sizes reflected in JJPX01.0061 (CFSAN003353) confirming the accuracy of inversion in the closed genome.

Despite having short-read Illumina sequences available at NCBI, this inversion went undetected in these strains. To our knowledge, this inversion is the largest ever identified in *Salmonella*, surpassing the next largest known inversion of 1.62 Mbp found in *Salmonella* Typhimurium (28). While it has been noted that serovar Typhimurium tends to invert portions of its genome around ribosomal RNA (rRNA) regions, those segments are comparatively small, measuring only 70-80 kb, unlike the inversion observed in these isolates (29, 30). Furthermore, the flanking regions of this inversion do not appear to contain rRNA; instead, they are associated with a Gifsy-1 phage, which appears to have become grounded due to its inability to excise as evidenced by the absence of the anti-repressor gene in the prophage (31). Additionally, while these isolates were all collected from environmental sources, it is possible that the genetic inversion coupled with the high sequence identity may be related to host-specific adaptations (27).

### Closely related *Salmonella* strains exhibit distinct pathogenicity dynamics in a live *C. elegans* host

The *Salmonella* strains tested shortened worm survival as compared to animals on OP50. Importantly, these strains showed differential impacts on survival rates with significantly different time to death (TD_50_) for each, with trends that were similar between two independent trials. As a positive control, we exposed *C. elegans* to *S*. Typhimurium CFSAN000741 and found that, as previously reported, it shortens *C. elegans* mean survival by 15%-30% in independent trials (Figure 3A, Table 2) (11, 32). Upon testing the 6 strains used in this study, we found that the strongest, consistent pathogenic impact was seen upon exposure to CFSAN003353, which caused a significantly greater lifespan reduction as compared to Typhimurium (53% v 15%) (Figure 3B, Table 2). CFSAN001469 and CFSAN00859 also caused greater survival diminution compared to CFSAN000741but their impacts were comparatively modest across trials (Table 2).

**Table 2:**
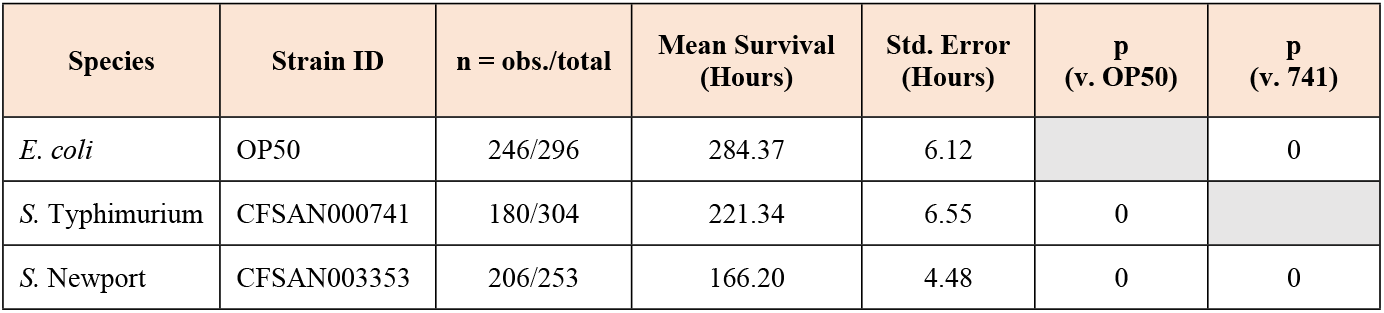
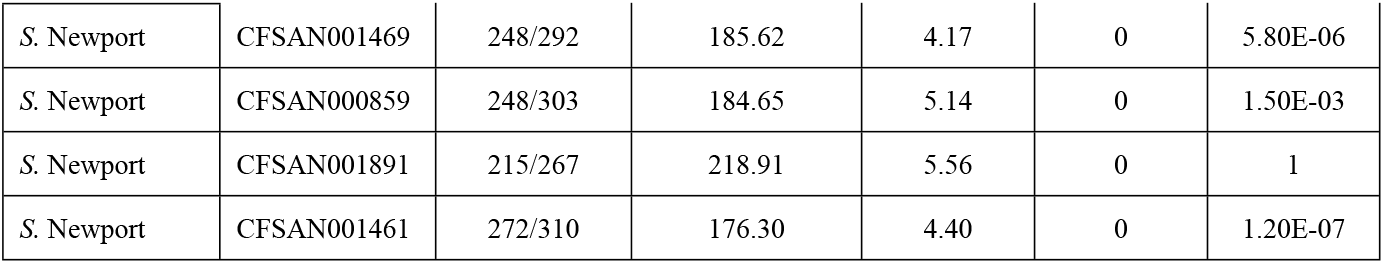
Impact of *Salmonella* strains on *C. elegans* Survival.

**Fig 3.**
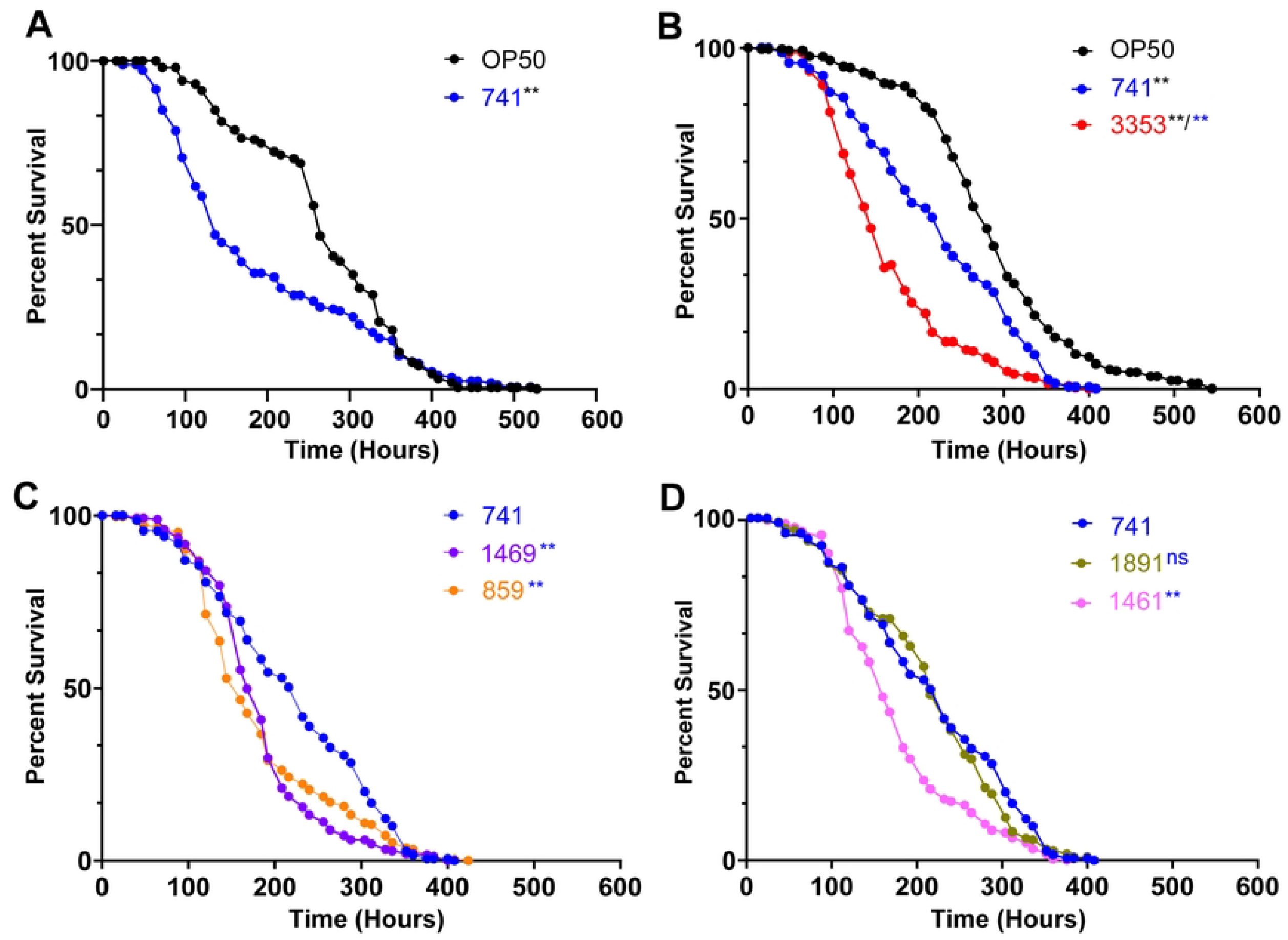
Impact of *Salmonella* strains on survival of *C. elegans*. **A:** *S*. Typhimurium (741, blue) shortens the survival of *C. elegans* adults compared to the normal diet of OP50 (black). **B:** *C. elegans* lifespan is shortened significantly more upon infection with CFSAN003353 (3353, red) compared to 741. **C:** CFSAN001469 (1469, purple) and CFSAN00859 (859, orange) cause greater reduction in worm lifespan compared to 741. **D:** Despite exhibiting the same PFGE pattern with the preserved inversion, strains CFSAN001461 (1461, pink) and CFSAN001891 (1891, olive) shorten worm lifespan to different degrees. 1461 causes significantly greater reduction than the control 741, whereas 1891 has the same magnitude of impact as 741. Data shown is combined from two independent trials (Table 2) and represents mean survival of the population in hours (m) ± SEM. P <0.0001 (**). ns = not statistically significant. Colors of asterisks indicate the strain being used for comparison.

Strikingly, we found that two of the strains, CFSAN001891 and CFSAN001461, which contain the large inversion and are > 99% identical to each other, exhibited distinctly different TD_50_s. In addition to the 15-30% shortening induced by CFSAN000741, CFSAN001461 exposure aggravated pathogen-driven survival diminution by an additional 25 – 30%, from ∼236 hours to ∼150 hours, whereas CFSAN001891 had the same effect as CFSAN000741 (Figure 3C, D, Table 2). Together, these experiments underscored the feasibility of using *C. elegans* to measure the virulence capacity of environmental *Salmonella* strains as well as nuanced pathogenicity differences resulting from genomic events.

## Conclusion

The *C. elegans* model has proven to be an invaluable tool for assessing a phenotype that might otherwise have gone unnoticed. Given that there were fewer than ten confirmed illnesses linked to Newport-1015 (<10 from 2000-2018), the virulence phenotype discovered in this study is surprising and suggests that this genomovar may not be preferentially adapted to human hosts. The isolates studied here offer a unique opportunity to elucidate the underlying regulatory mechanisms that may contribute to the differences in virulence among *Salmonella* serovars.

Moreover, due to the highly conserved nature of the genomes, distinguishing differences between the two genomovars (Newport-61 vs. Newport-1015) can currently only be achieved using PFGE or long-read sequencing. Comparative genomics alone is insufficient in drawing conclusions about the virulence potential of closely related strains of *Salmonella*. For this reason, expression studies will be required to identify the mechanisms underlying these phenotypes, along with genome-wide screening to understand the roles that specific genes play in the infection process.

## Acknowledgements

This work was supported by National Institutes of Health grants R01AI176326, R21AG083329 and K07AG078287 to AG. Thanks to Hannah Kang for help provided with the worm experiments.

## Notes

### Competing Interest Statement

The authors have declared no competing interest.

